# A stuttering-associated *Gnptab* variant alters fine-motor kinematics

**DOI:** 10.64898/2025.12.08.693031

**Authors:** Devin Bishop, Siwei Wang, Shahriar SheikhBahaei

**Affiliations:** Department of Neurobiology & Behavior, Stony Brook University, Stony Brook, NY 11794, USA; Center for Nervous System Disorders, Stony Brook University, Stony Brook, NY 11794, USA; Department of Neurobiology & Behavior, Stony Brook University Renaissance School of Medicine, Stony Brook, NY 11794, USA

**Keywords:** basal ganglia, cortex, fine motor behavior, Gnptab, grasping

## Abstract

Precise reach-to-grasp movements rely on complex cortico-basal ganglia and cerebellar circuits, and can be affected by cellular pathway defects. We investigated whether mild *Gnptab* deficiency, affecting mannose–6–phosphate–dependent lysosomal enzyme targeting, impairs fine motor behaviors in mice. Using the staircase test, high-frame-rate video, and markerless tracking, we assessed reaching and grasping in *Gnptab*-mutant and control littermates. Our data suggest that *Gnptab*-mutant mice retrieved about 75% fewer pellets than controls and exhibited abnormal grasping, characterized by increased wrist extension and larger digit angles, decreased digital velocity, and shorter reach distances, while elbow angles remained largely unchanged. These findings suggest a specific deficit in grasp shaping and trajectory control, rather than overall limb positioning. Our data establishes a quantitative link between *GNPTAB*-related lysosomal pathway disruption and fine-motor impairments, providing a valuable model for understanding how cellular dysfunction impacts motor circuit function.

## Introduction

Precise control of fine motor movements is essential for behaviors such as reaching, grasping, and object manipulation (Olivier et al. 2007). These actions rely on coordinated activity across cortico-striatal, cerebellar, and brainstem motor circuits that integrate proprioceptive and visual feedback to guide movement timing and accuracy (Asan, McIntosh, and Carmel 2021; Wolpert, Ghahramani, and Jordan 1995; Scott 2004). Developmental or acquired disruptions in these circuits can impair coordination and precision, leading to deficits in both motor and communicative behaviors. Understanding the genetic and cellular mechanisms that govern fine motor control is therefore critical for identifying pathways that contribute to neuromotor and neurodevelopmental disorders.

Mutations in *GNPTAB*, which encodes the α/β subunits of N-acetylglucosamine-1-phosphotransferase, disrupt the tagging of lysosomal enzymes with mannose-6-phosphate and lead to mislocalization of hydrolases outside the lysosome, causing lysosomal storage disorders (Velho et al. 2019). Specific, milder mutations in GNPTAB do not cause lysosomal storage disorders but have been linked to developmental stuttering and other speech-motor deficits (Kang et al. 2010). Knock-in *Gnptab* mice exhibit phenotypes that parallel aspects of these disorders, including atypical ultrasonic vocalizations, disrupted sensorimotor control, and altered astrocyte morphology (Adeck et al. 2024; Millwater, Bragg, et al. 2025; Millwater, Weinhold, et al. 2025). However, while previous studies have focused primarily on gross motor and vocal behaviors (Millwater, Weinhold, et al. 2025), the effects of *Gnptab* deficiency on fine motor coordination remain unknown.

In this study, we employed behavioral testing and kinematic analysis of pellet-reaching tasks to investigate fine motor control in *Gnptab*-mutant mice and their control littermates. We hypothesized that *Gnptab* deficiency would impair reaching performance, reflected by decreased grasp success and altered joint dynamics during pellet retrieval. Using the staircase test (Baird, Meldrum, and Dunnett 2001), high-frame-rate video recording, and markerless pose estimation (Mathis et al. 2018), we quantified forelimb kinematics, including velocity profiles and joint-angle coordination. Together, our data provide a quantitative framework for understanding how disruptions in the *GNPTAB*-dependent cellular transport system impact fine motor behavior and may offer insight into the neural mechanisms linking lysosomal dysfunction to developmental motor coordination deficits.

## Methods

### Animals

Male *Gnptab*-mutant mice and their wild-type littermates, aged 6–8 weeks, were used for all experiments. Mice were maintained on a C57BL/6 background and housed in a temperature-controlled facility (24°C) with a 12-hour light/dark cycle and *ad libitum* access to food and water. All procedures were approved by the Institutional Animal Care and Use Committee and conducted in accordance with NIH guidelines for animal research.

### Staircase Test Apparatus and Procedure

The staircase test (**Figure 1A**) was conducted using a standard apparatus from Campden Instruments consisting of a clear Perspex box with a central platform and two staircases containing baited steps (Baird, Meldrum, and Dunnett 2001). The 4^th^, 5^th^, and 7^th^ steps were loaded with one or two 20 mg precision pellets (Bio-Serv, NJ, USA) to assess reaching and grasping performance. *Gnptab*-mutant and control mice were placed individually in the apparatus for a single one-hour session. The number of pellets displaced and retrieved from each step was recorded as a measure of grasping performance. All testing was conducted during the light phase (13:00–17:00 h) to control for circadian effects.

**Figure 1.**
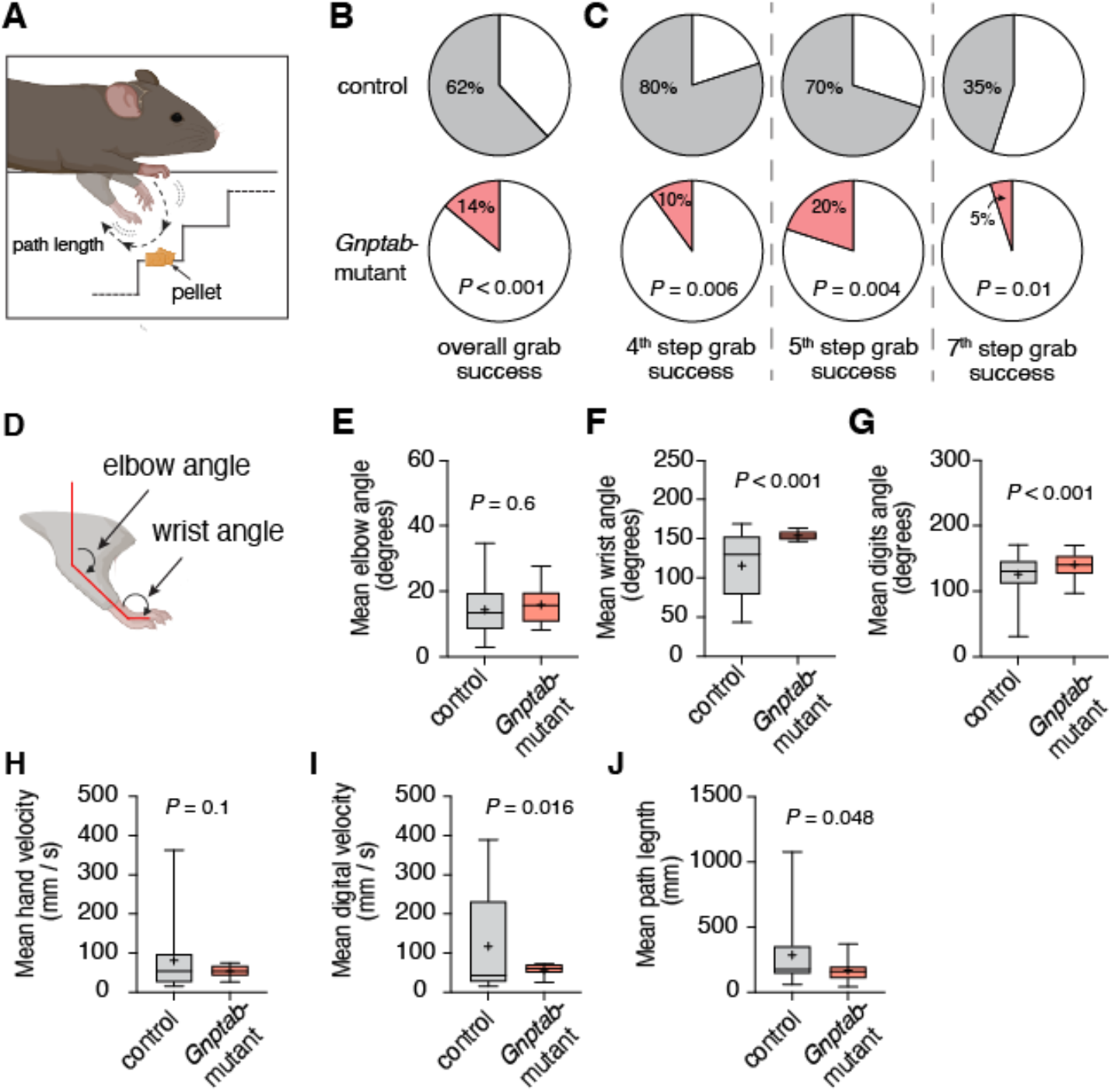
*Gnptab* mutation reduces pellet retrieval and alters grasp kinematics in mice. (**A**) Schematic of the staircase reaching setup illustrating pellet positions and the measured reach trajectory. (**B**) Overall, the grab success across all steps is reduced in *Gnptab*-mutant mice (control: 62%; *Gnptab*-mutant: 14%). (**C**) Success rates between steps show deficits at individual stair positions. (**D**) Diagram showing joint angles used for kinematic analysis. (**E–G**) Mean elbow angle was unchanged, whereas wrist angle and digit angle were larger in *Gnptab*-mutant mice, reflecting a more extended wrist and open grasp posture during pellet approach. (**H–I**) Mean hand velocity was not different, while mean digital velocity was reduced in *Gnptab*-mutant mice. (**J**) Reach trajectories were shorter in *Gnptab*-mutant mice. *P* values – B&C: Fisher’s exact test; E-J: unpaired *t*-test. s – seconds.

### Video Recording and DeepLabCut Analysis

Mouse behavior during the staircase test was recorded using high-resolution Basler acA1300 (Basler AG, Germany) cameras positioned perpendicular to the staircase to capture optimal views of the reaching and grasping movements. Videos were recorded at 60 frames per second (fps) and saved in MP4 format for subsequent analysis. Markerless pose estimation was performed using DeepLabCut (Mathis et al. 2018). Approximately 1400 frames from representative videos were manually labeled for key points, including the snout, shoulder, elbow, wrist, digits, and reference points for food pellets on the staircase. A ResNet-50–based DLC network was trained on diverse frames to ensure generalization, achieving test error < 10 pixels. The trained network tracked body parts across all videos, and the resulting x,y coordinates were analyzed using DLC2Kinematics (Mathis et al. 2020). This pipeline calculated kinematic parameters, including velocity profiles and joint angles.

### Statistical Analysis

Data are presented as mean ± SEM. Comparisons between *Gnptab*-mutant and control mice were conducted using unpaired *t*-tests or the Fisher’s exact test, as appropriate.

## Results

### Pellet Retrieval Performance

We first investigated pellet retrieval in *Gnptab*-mutant and control mice. Control mice successfully retrieved an average of 1.3 ± 0.6 pellets per hour, whereas *Gnptab*-mutant mice retrieved only 0.3 ± 0.4 pellets (**Figure 1A-B**). This represents a ∼ 75 % reduction in retrieval success (*P* < 0.001, Fisher’s exact test). Deficits were consistent across steps, with reduced success observed on steps 4 (*P* = 0.006, Fisher’s exact test), 5 (*P* = 0.004, Fisher’s exact test), and 7 (*P* = 0.010, Fisher’s exact test; **Figure 1C**). These data demonstrate a robust impairment in fine-motor reaching performance in *Gnptab*-mutant mice.

### Kinematic Analysis

Computer vision-based kinematic analysis revealed distinct alterations in limb and digit movement patterns between groups (**Figure 1D-J**). Elbow joint angles during grasping did not differ between *Gnptab*-mutant and control mice (14 ± 1° vs 16 ± 2°, *P* = 0.6, *t*-test; **Figure 1E**), suggesting gross limb positioning was largely preserved. However, wrist angles were significantly more extended in *Gnptab*-mutant mice (154 ± 2° vs 116 ± 7° in control mice, *P* < 0.001, *t*-test; **Figure 1F**), and the mean digit angle was also greater (*P* < 0.001, *t*-test; **Figure 1G**), indicating an abnormally open palm posture. Mean hand velocity was reduced in mutants (53 ± 6 mm/s vs 81 ± 15 mm/s in control mice, *P* = 0.1, *t*-test; **Figure 1H**). Mean digital velocity was also significantly reduced (*P* = 0.02, *t*-test; **Figure 1I**). The coefficient of velocity variation was lower in *Gnptab*-mutant mice (177 ± 11 vs 246 ± 24 in control), suggesting reduced adaptive modulation of movement speed across trials. Path length was also shorter (168 ± 34 mm vs 286 ± 46 mm in control mice; *P* = 0.05, *t*-test; **Figure 1A & J**), indicating truncated or incomplete reach trajectories. Together, these results suggest that while gross limb positioning was maintained, *Gnptab*-mutant mice exhibited abnormal wrist extension, reduced velocity variability, and shortened reach trajectories, consistent with impaired grasp formation and fine-motor control (Mathis et al. 2018; Bishop et al. 2022; Skrobot et al. 2024).

## Discussion

This study demonstrates that mild *Gnptab* deficiency disrupts fine-motor performance during a reaching and grasping task. In our experimental setup (**Figure 1A**), *Gnptab*-mutant mice retrieved fewer pellets and showed slower, less variable limb movements, characterized by extended wrist and finger postures. These findings indicate a disruption in the fine motor control mechanisms underlying reach-to-grasp behavior, consistent with broader sensorimotor coordination deficits that were previously described in these animals (Millwater, Weinhold, et al. 2025) and in people who stutter (Falk, Müller, and Dalla Bella 2015).

*GNPTAB* encodes the α/β subunits of N-acetylglucosamine-1-phosphotransferase, the enzyme responsible for tagging lysosomal hydrolases with mannose-6-phosphate (Velho et al. 2019). Disruption of this pathway leads to lysosomal enzyme mislocalization and cellular stress, which can impact multiple brain regions involved in goal-directed movement. The observed motor impairments may therefore arise from altered signaling within cortico-cerebellar and cortico-striatal (Maguire, Yoo, and SheikhBahaei 2021; Turk et al. 2021; Chang and Guenther 2019; Yang et al. 2016; Craig-McQuaide et al. 2014) pathways that integrate sensory feedback to refine ongoing actions (Sitek et al. 2016; Matsuhashi et al. 2023; Watkins et al. 2008). Impaired communication within these circuits could underlie the reduced reach precision and abnormal grasp closure observed in *Gnptab*-mutant mice.

Previous behavioral observations have shown that *Gnptab* mutations alter both vocal and gross motor behaviors in *Gnptab*-mutant mice (Han et al. 2019; Millwater, Bragg, et al. 2025; Millwater, Weinhold, et al. 2025). Our data provide a direct link between *Gnptab* deficiency and measurable impairments in fine motor coordination, connecting the cellular lysosomal dysfunction to circuits that control fine-motor mechanisms within distributed motor circuits.

The combination of kinematic abnormalities suggests a distributed network effect involving cortico-basal ganglia-thalamo-cortical circuits. Disruption of these circuits has also been proposed in developmental stuttering, a speech-motor timing disorder (Turk et al. 2021; Chang and Guenther 2019; Alm 2004), supporting the idea that *GNPTAB*-related deficits in motor timing may extend across limb and orofacial systems. Future work should integrate behavioral and kinematic measures with neural recordings to determine the circuit-level mechanisms underlying these deficits. Including both sexes, testing additional paradigms such as single-pellet reaching or ladder-rung walking, and applying longitudinal designs will help further define *GNPTAB*-related motor dysfunction.

The putative cortical-basal ganglia and cortico-cerebellar circuits we propose as vulnerable to *Gnptab* mutations in rodents have close homologs in the human speech motor system (Bostan and Strick 2018; Simonyan 2014; Chang and Guenther 2019; Foster et al. 2021). Convergent models and imaging have demonstrated that these circuits support the initiation, sequencing, stabilization, and integration of rapid orofacial movements, as well as the integration of sensory feedback with forward predictions (Tourville and Guenther 2011; G et al. 2022). Reports of atypical activity and connectivity within these networks in individuals who stutter—together with subtle deficits on non-speech fine-motor tasks—support a domain-general motor-control instability rather than a purely speech-specific anomaly (Gracco, Sares, and Koirala 2022; Shojaeilangari et al. 2021; Yang et al. 2016; Sitek et al. 2016; Smits-Bandstra, De Nil, and Saint-Cyr 2006; Hajj et al. 2023; Höbler et al. 2022; Mersov et al. 2016). Our data add a cellular mechanism to this framework: lysosomal-pathway dysfunction within glia–neuron networks can perturb motor timing and coordination at the circuit level, consistent with evidence that lysosomal defects disrupt synaptic and network function (Gorelik et al. 2022; SheikhBahaei, Millwater, and Maguire 2023; Adeck et al. 2024; Parenti, Medina, and Ballabio 2021).

Considering that the studied mutation in *GNPTAB* is linked to stuttering in humans, we propose that a subset of stuttering disorders may be due to vulnerabilities in conserved motor control circuits. This aligns with human genetic links to disruptions in the lysosomal enzyme-targeting pathway and with mouse models carrying human GNPTAB mutations that mimic stuttering-like vocalization issues (Kang et al. 2010; Han et al. 2019; Barnes et al. 2016; Millwater, Bragg, et al. 2025; Millwater, Weinhold, et al. 2025). This account predicts heterogeneity aligned with circuit properties, such as variability and connectivity within cortico-basal ganglia-thalamo-cortical circuits, which are already linked to stuttering disorders in humans (SheikhBahaei, Millwater, and Maguire 2023; Neef et al. 2018; Cai et al. 2014; Neef and Chang 2024; Chang et al. 2025). In this regard, the *Gnptab*-mutant mouse model provides a practical platform for connecting cellular lysosomal defects to these circuit-level indicators.

## Acknowledgements

We thank the NIMH Rodent Behavior Core for their assistance with the pilot study and Julianna Saxena for their comments on an earlier version of the manuscript. This work was supported, in part, by the MH002952 and NS009420.

